# Phosphodiesterase 5 inhibition improves contractile function and restores transverse tubule loss and catecholamine responsiveness in heart failure

**DOI:** 10.1101/248187

**Authors:** Michael Lawless, Jessica L. Caldwell, Emma J. Radcliffe, Charlotte E.R. Smith, George W.P. Madders, David C. Hutchings, Lori S. Woods, Stephanie J. Church, Richard D. Unwin, Graeme J. Kirkwood, Lorenz K. Becker, Charles M. Pearman, Rebecca F. Taylor, David A. Eisner, Katharine M. Dibb, Andrew. W Trafford

**Affiliations:** Division of Cardiovascular Sciences, Unit of Cardiac Physiology, School of Medical Sciences, Faculty of Biology, Medicine and Health, The University of Manchester, Manchester Academic Health Science Centre, Manchester Academic Health Science Centre, 3.24 Core Technology Facility, 46 Grafton Street, Manchester, M13 9NT, United Kingdom.; Division of Cardiovascular Sciences, Centre for Advanced Discovery and Experimental Therapeutics), School of Medical Sciences, Faculty of Biology, Medicine and Health, The University of Manchester, Manchester Academic Health Science Centre, Manchester Academic Health Science Centre, 3.24 Core Technology Facility, 46 Grafton Street, Manchester, M13 9NT, United Kingdom.

## Abstract

Heart failure (HF) is characterized by poor survival, a loss of catecholamine reserve and cellular structural remodeling in the form of disorganization and loss of the transverse tubule network. Indeed, survival rates for HF are worse than many common cancers and have not improved over time. Tadalafil is a clinically relevant drug that blocks phosphodiesterase 5 with high specificity and is used to treat erectile dysfunction. Using a sheep model of advanced HF, we show that tadalafil treatment improves contractile function, reverses transverse tubule loss and restores calcium transient amplitude and the hearts response to catecholamines. Accompanying these effects tadalafil treatment normalized BNP mRNA and prevented development of subjective signs of HF. These effects are independent of changes in myocardial cGMP content and are associated with upregulation of both monomeric and dimerized forms of protein kinase G and of the cGMP hydrolyzing phosphodiesterase 2 and 3. We propose that the molecular switch for the loss of transverse tubules in HF and their restoration following tadalafil treatment involves the BAR domain protein Amphiphysin II (BIN1) and the restoration of catecholamine sensitivity is through reductions in G-protein receptor kinase 2, protein phosphatase 1 and protein phosphatase 2A abundance following phosphodiesterase 5 inhibition.

---

Impaired responsiveness to catecholamines is a hallmark of the failing heart and is detectable both *in vivo* and in isolated ventricular myocytes ^1,2^. The mechanisms responsible for the attenuated catecholamine effects in heart failure (HF) are varied and include reduced adenylate cyclase activity and enhanced G protein receptor kinase (GRK2) and intracellular protein phosphatase activity (PP1 and PP2A) which together lead to a decrease in cAMP dependent signaling and impaired PKA-dependent target phosphorylation ^1,3,4^. Given the functional distribution of β-adrenergic receptors (β-ARs) and G-proteins across the surface sarcolemma and transverse tubule (TT) membrane ^5–7^ a further factor proposed to contribute to impairment of the β-adrenergic signaling cascade in HF is the reduction of transverse tubule (TT) density seen in many pre-clinical models and human HF ^8–11^.

In addition to the classical cAMP-dependent process, the myocardial response to catecholamine stimulation is also regulated by the cGMP-PKG signaling axis consisting of the β_3_-AR / soluble guanylate cyclase (sGC) and natriuretic peptide / particulate guanylate cyclase (pGC) pathways (reviewed by Tsai and Kass ^12^). The outcome of cGMP dependent activation depends on the source of activating cGMP; that activated by sGC inhibiting the β-AR response and, pGC derived cGMP having no effect ^13,14^. Beyond the role of PKG, GRK2 and protein phosphatases in determining the outcome of β-AR stimulation, the intracellular pools of cAMP and cGMP are also differentially regulated by phosphodiesterases (PDEs) suggesting highly compartmentalized regulation of the cyclic nucleotides and thus catecholamine responsiveness of the healthy ventricular myocardium e.g. ^15–18^.

Given the negative impact of acute PDE5 inhibition on the inotropic and lusitropic response to catecholamines and the established loss of catecholamine reserve in HF it is somewhat surprising that an emerging body of evidence suggests PDE5 inhibition is clinically cardioprotective in type II diabetes ^19^, left ventricular hypertrophy ^20^ and in patients with HF with reduced ejection fraction (systolic HF) ^21^. Similarly, in experimental models, PDE5 inhibition shows cardioprotective effects in pulmonary hypertension ^22^, myocardial infarction ^23–26^ and following aortic banding ^27^. However, in each of these cases PDE inhibition was commenced either before or given concurrently with the disease intervention. Such an experimental approach complicates interpretation of whether the intervention is therapeutically useful in a setting of established disease or is acting by preventing disease development. In most ^28,29^, but not all ^30^ experimental studies where PDE5 inhibition has been commenced once some degree of left ventricular remodeling has occurred the findings remain supportive of a cardioprotective effect. However, in the ‘positive’ studies the extent of disease progression to symptomatic HF is unclear and data on survival outcomes is generally missing.

Given these considerations the hypothesis examined is that phosphodiesterase 5 inhibition is beneficial in systolic HF through restoration of catecholamine responsiveness. As such, the primary aim of the present study was to determine if PDE5 inhibitor treatment, instigated at an advanced disease stage once contractile dysfunction and attenuated catecholamine responsiveness are established, is capable of reversing these effects. The secondary aim of the study was to determine if changes in contractile and catecholamine responsiveness were associated with structural (TT) remodeling and to elucidate the underlying molecular mechanisms of such TT remodeling. The final aim of the study was to determine the underlying mechanisms that contribute to the restoration of catecholamine responsiveness.

The major findings are that PDE5 inhibition with tadalafil restored catecholamine responsiveness and partially reversed contractile *in vivo*. At the cellular level, PDE5 inhibition restored the amplitude of the systolic calcium transient to control levels. These changes were associated with a restoration of TT density but β-AR abundance (β_1_ and β_2_) remained unaltered in HF and following tadalafil treatment. However, downstream regulators of catecholamine signalling including GRK2, PKG, PDE2, PDE3, PP1 and PP2A were all differentially regulated by HF and PDE5 inhibition. Our findings also implicate the BAR domain protein amphiphysin II (AmpII *aka* BIN1) as a key driver of the TT changes seen in response to HF and PDE5 inhibitor treatment. Additionally, we found that tadalafil treatment reversed myocardial changes in BNP expression and that this was associated with the prevention of the development of subjective HF symptoms.

## Results

### PDE5 inhibition improves cardiac contractility in vivo and systolic calcium in vitro

We first sought to determine if tadalafil treatment was associated with any changes in cardiac contractility or cardiac dimensions. In line with our previous findings ^1,31–33^ left ventricular dilatation measured at end diastole (end diastolic internal dimension; EDID) and the end of the systolic phase (end systolic internal dimension; ESID) as well as free wall thinning were observed following tachypacing. In a subset of animals where echocardiographic assessments were repeated every week, tadalafil treatment did not reverse left ventricular dilatation (Figure 1A, all values *p* < 0.01 vs pre-pacing). Similarly, systolic wall thinning occurred in tachypaced animals; an effect not reversed by tadalafil treatment (*p* < 0.05 vs pre-pacing). However, after three weeks of tadalafil treatment diastolic wall thickness was indistinguishable from pre-pacing levels (Figure 1B; *p* = 0.09). In sheep the cardiac apex is positioned over the sternum and it is difficult to obtain four chamber apical views in adult animals. Therefore, we used the short axis fractional area change measured at mid papillary muscle level as a surrogate for ejection fraction. Following 4-weeks of tachypacing fractional area change was reduced from pre-pacing values by 43 ± 7 % (Figure 1C) in the untreated group and to the same extent in the tadalafil group before treatment commenced (43 ± 3%, both groups *p* < 0.001 vs pre-pacing values). However, tadalafil treatment increased fractional area change such that by the end of the study fractional area change was 16 ± 8 % greater than at 4-weeks (*p* < 0.05) and 35 ± 24 % greater than in the end-stage untreated group (*p* < 0.01). In agreement with the *in vivo* contractility findings and our previous study ^1^, the amplitude of the systolic calcium transient was reduced by 66 ± 14 % in HF (Figure 1D & E, *p* < 0.05). Importantly, the reduction in calcium transient amplitude was established by 4 weeks of tachypacing (by 74 ± 8 %, *p* < 0.05). However, following tadalafil treatment the amplitude of the systolic calcium transient was 399 ± 150 % of HF values (*p* < 0.001) and indistinguishable from control (132 ± 32 % of control; *p* = 0.49). Therefore, tadalafil treatment restored systolic calcium to a greater extent than *in vivo* contractility. Whilst tadalafil treatment augmented cardiac contractility and systolic calcium, the additive effects of tadalafil treatment on blood pressure were minimal. Previously we have reported that systolic, diastolic and mean blood pressure decrease in HF ^34^; an observation repeated here (Table 1). However, tadalafil treatment had no further effect on blood pressure which was indistinguishable from both the 4-week tachypaced and HF groups. Thus, the changes in cardiac function associated with tadalafil treatment are not due to changes in peripheral blood pressure.

**Table 1.**
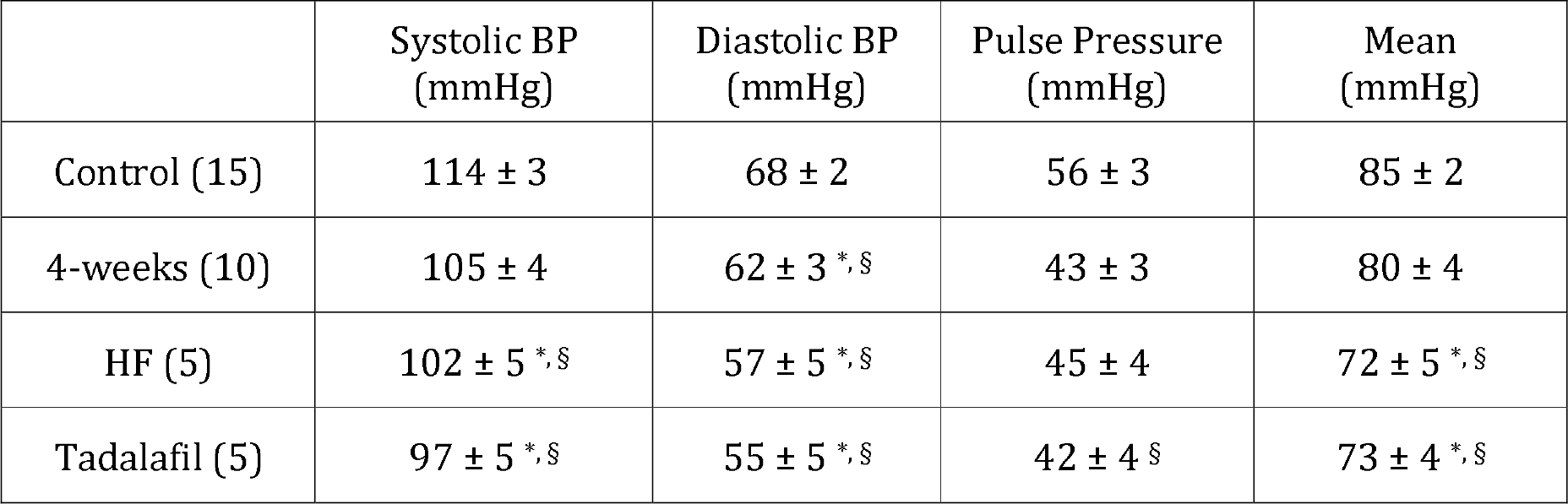
The influence of HF and tadalafil treatment on blood pressure measured by tail cuff plethysmography. Summary blood pressure data (mean ± standard error). Values in brackets denote number of animals in each condition. *, denotes *p* < 0.05 vs. control (One way ANOVA with SNK correction for multiple comparisons). ^§^, denotes *p* < 0.05 vs. control (paired t-test).

**Figure 1.**
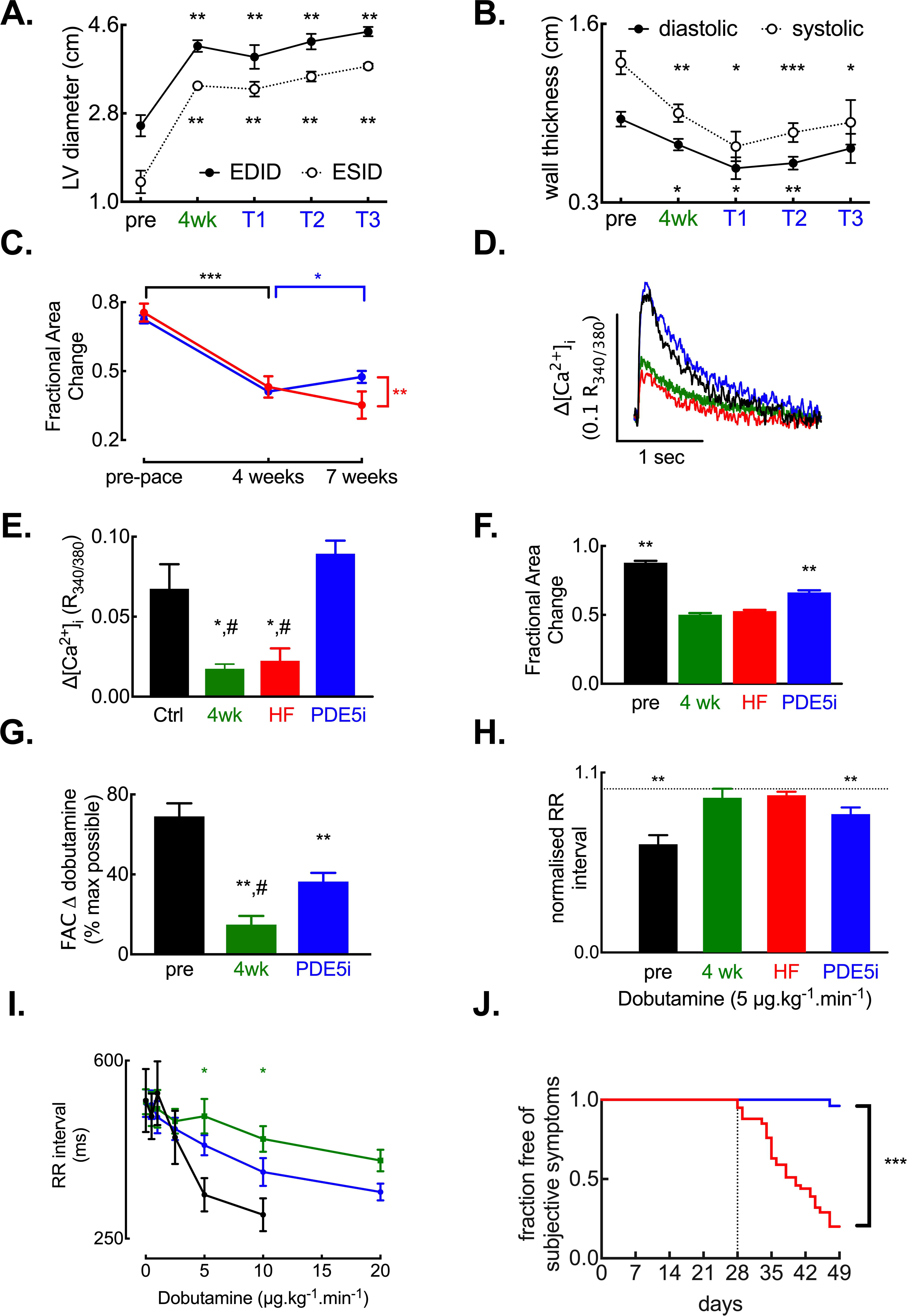
Tadalafil treatment restores indices of cardiac function in heart failure. **A**. Changes in left ventricular dimensions with tachypacing and response to tadalafil treatment (paired data, *N* = 4 each time point; **, *p* < 0.01 by RM-ANOVA). ‘pre’ denotes before commencement of tachypacing, ‘4wk’ denotes 4-weeks of tachypacing and T1, T2 and T3 denote 1, 2 & 3 weeks of tadalafil treatment. **B**. Changes in left ventricular free wall thickness in response to tachypacing and tadalafil treatment (paired data, *N* = 4 each time point; *, **, *** respectively *p* < 0.05, 0.01 & 0.001 by RM-ANOVA). **C**. Echocardiographic assessment showing short-axis fractional area change in the absence of exogenous catecholamine stimulation: *, *p* <0.05 vs. 4-week time point (tadalafil arm); **, *p* < 0.01 vs. HF group at 7 weeks; ***, *p* < 0.001 vs. pre-pace function (both groups). Statistics by mixed models analysis. *N* = HF, 5 at all time points; tadalafil, 20 at pre-pace, 19 at 4-weeks and 18 at 7 weeks. **D**. Representative systolic calcium transients for control (black), 4-week paced (green), HF (red) and tadalafil (blue) groups. **E**. Summary data for systolic calcium transients; *, *p* < 0.05 vs. control; #, *p* < 0.05 vs. tadalafil. (*n*/*N* = 34/16 control, 6/2 following 4 weeks tachypacing, 7/3 HF, 24/6 Tadalafil). **F**. Summary data of echocardiographic assessments following 5 µg/kg/min dobutamine infusion: **, *p* < 0.01 vs. HF by one-way ANOVA. *N* = pre-pacing, 4; 4-weeks, 3; HF, 4; tadalafil, 3. **G**. Summary data showing dobutamine induced change in fractional area change derived from paired (repeated measures) data in panel F. Data is expressed as a percentage of the maximum possible response in each group. **H**. Summary data showing RR intervals normalized to pre-dobutamine (5 µg.kg/min) RR interval: **, *p* < 0.01 vs. pre-dobutamine RR interval by t-test. *N* = pre-pacing, 5; 4-weeks, 5; HF, 6; tadalafil, 5. **I**. Paired data from 5 animals showing RR interval – dobutamine dose response relationship. *, *p* < 0.05 by mixed models analysis. **J**. Kaplan-Meier plot showing fraction of animals free from subjectively assessed signs of HF (*N* = HF, 42; tadalafil, 27). Tadalafil treatment commenced 28 days after tachypacing (dashed line): ***, denotes *p* < 0.001 (by Cox regression-based test).

### PDE5 inhibition restores the inotropic and chronotropic response to catecholamine stimulation

Since HF is associated with a reduced responsiveness to β-adrenergic stimulation ^1,31^ and impaired catecholamine responses are associated with reduced survival in HF patients ^35^ we next determined if tadalafil treatment was also associated with restoration of catecholamine responsiveness. We first examined cardiac contractile responses to intravenous infusion of the exogenous β-adrenergic agonist dobutamine (Figure 1F & G). In paired samples (pre-pacing, 4-weeks and tadalafil treated), there was a greater response to dobutamine infusion after tadalafil treatment than after 4-weeks of tachypacing (Figure 1G) indicating that chronic PDE5 inhibition in HF augments catecholamine sensitivity. As we have reported previously in a separate study ^31^, HF attenuated the chronotropic response to dobutamine. In pre-paced animals, 5 µg/kg/min dobutamine increased heart rate and shortened RR interval by 36 ± 10 % (Figure 1H; *p* < 0.01) yet had no effect on RR interval following 4-weeks of tachypacing or in end-stage HF (the change in RR interval being 5.4 ± 8 % and 3.8 ± 2 % respectively). Conversely, at this infusion rate following tadalafil treatment, heart rate increased resulting in RR interval shortening by 26 ± 6 % (*p* < 0.01). To further understand the attenuated chronotropic response, we also investigated the dose-response to dobutamine in a cohort of animals at three time points; pre-pacing, 4-weeks of tachypacing (but before tadalafil treatment commenced) and following 3 weeks of tadalafil treatment (Figure 1I). It is clear that the sensitivity to dobutamine was reduced at 4-weeks of tachypacing and restored toward control values following tadalafil treatment. Assuming that the maximal achievable heart rate is equal to that in pre-paced animals treated with 10 µg/kg/min dobutamine, the calculated EC50 concentration of dobutamine is 3.6, 34.4 and 10.7 µg/kg/min in the pre-pacing, 4-weeks tachypaced and tadalafil treated groups respectively (*p* < 0.005).

To conclude the *in vivo* series of experiments, we sought to determine if tadalafil treatment was associated with any improvement in the subjective signs of HF (including lethargy, dyspnoea, cough or loss of bodily condition). In the untreated cohort, 34 of 42 animals developed at least two subjective signs of HF before the designated end of the study (49 days of tachypacing; all 34 animals were dyspnoeic at rest and coughed when restrained in a supine position) and were therefore humanely killed for subsequent in vitro experiments. Conversely, of the 27 animals randomly assigned to the tadalafil treatment group, all remained free of signs of HF to the completion of the study (day 49, Figure 1J). One tadalafil treated animal was found dead on day 19 of treatment; notably there were no prior subjective signs of HF and on post-mortem examination the pacing lead was found to have perforated the right ventricular wall and pulmonary congestion was not evident. Thus, compared to untreated animals, tadalafil treatment yielded a Cox’s proportional hazard ratio of 0.026 ± 0.03 (95 % CI, 0.003 – 0.19; *p* < 0.001). Additionally, there was no difference in the initial body weight between the 42 untreated (32.0 ± 1.1 kg) and 26 tadalafil treated animals (31.6 ± 1.3 kg) and, as such, body weight was not a significant covariate (*p* = 0.44). Finally, in a subset of the tadalafil treated HF animals (*N* = 13) the total plasma tadalafil content determined by mass spectrometry was 269 ± 39 µg/l and plasma tadalafil protein binding measured as 93 ± 0.6 % which is in good agreement with that in healthy human subjects ^36^. Thus, the lowest free tadalafil concentration in HF plasma following 3 weeks of daily dosing (20 mg daily) was 51 ± 8 nmol/l.

In addition to improving several *in vivo* measures of cardiac function and catecholamine sensitivity, chronic PDE5 inhibition also normalized the tachypacing induced increases in myocardial NPPB (B-type natriuretic peptide) mRNA (Figure 2A). Thus, tadalafil treatment at least partially restores both the inotropic and chronotropic response to catecholamines that are lost in HF and, in further agreement with the improvement in the subjective assessments of HF development, tadalafil treatment also reverses the increase in mRNA abundance of the widely accepted marker of HF, B-type natriuretic peptide.

**Figure 2.**
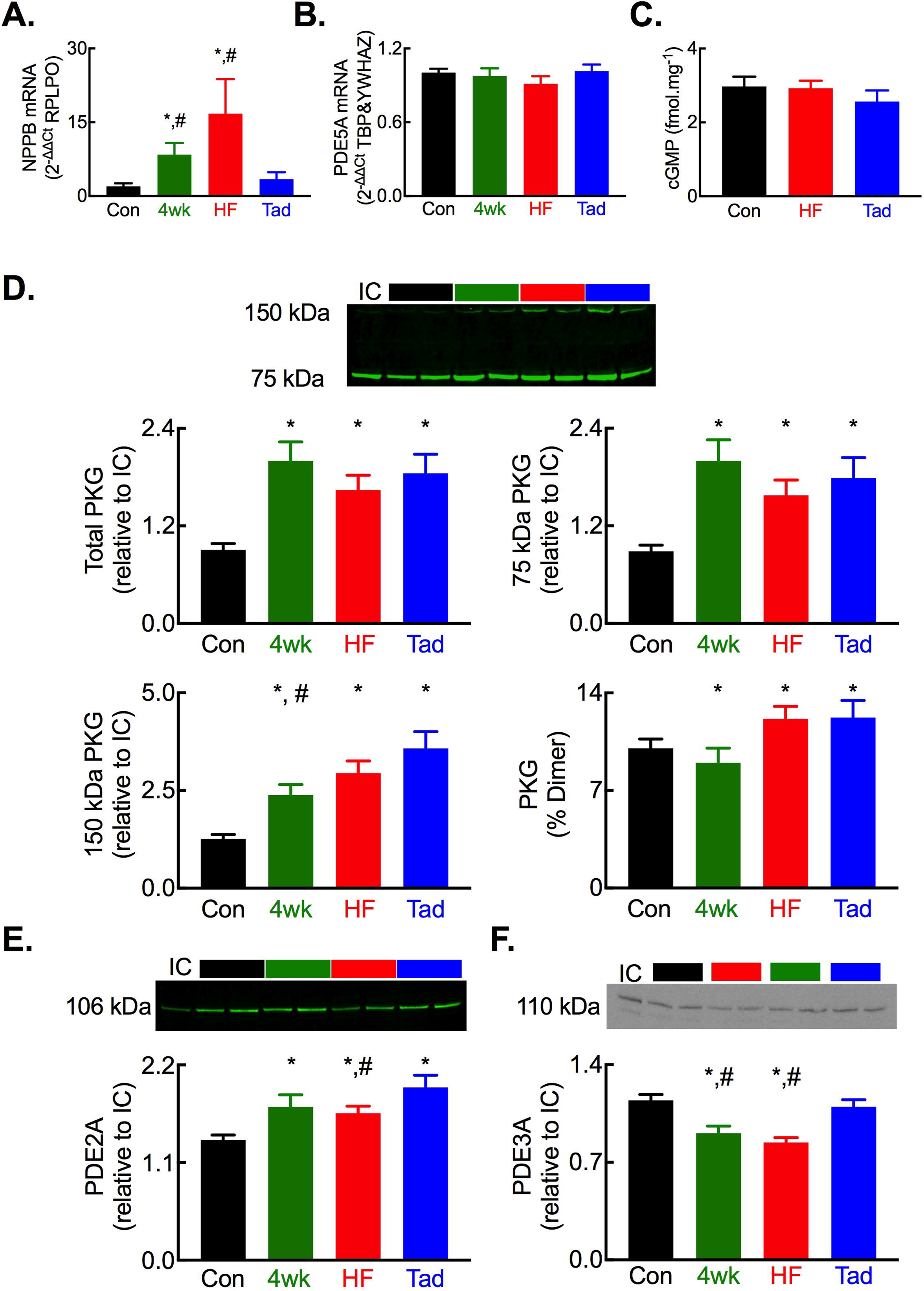
Altered abundance of regulators of cyclic GMP in HF and following tadalafil treatment. **A**. Summary quantitative PCR assessment of myocardial brain natriuretic peptide (NPPB) mRNA abundance normalized to the housekeeper RPLPO. **B**. Summary quantitative PCR assessment of myocardial phosphodiesterase 5A (PDE5A) mRNA abundance normalized to the housekeepers TBP and YWHAZ. **C**. Summary data showing no change in myocardial cGMP content. **D**. Changes in protein kinase G abundance assessed by non-denaturing PAGE showing (top) a representative blot, (middle) quantification of total and 75 kDa PKG and, (bottom) quantification of 150 kDa PKG and % total PKG as dimerized form. **E**. Altered PDE2A protein abundance showing (top) representative blot and, (bottom) summary data. **F**. Altered PDE2A protein abundance showing (top) representative blot and, (bottom) summary data. For all panels *, *p* < 0.05 vs control; #, *p* < 0.05 vs tadalafil. *N* = 8 each group.

### The impact of PDE5 inhibition on regulators of cGMP and the β-adrenergic signalling cascade

There was no change in the abundance of PDE5A mRNA (Figure 2B) or cGMP levels in myocardial homogenates (Figure 2C) in HF or following tadalafil treatment. Whilst unaltered PDE5 mRNA levels in HF have been reported previously ^29,37^, the unchanged myocardial cGMP content following tadalafil treatment was initially surprising. However, examining known regulators of cGMP in HF and following tadalafil treatment we found an increase in total, monomeric (75 kDa) and dimerized (150 kDa) PKG (Figure 2D). Total and 75 kDa PKG abundances were similar between the 4-week paced, HF and tadalafil groups. However, for the 150 kDa dimer of PKG, abundance was greater in the tadalafil group compared to the 4-week paced group and trended to an increase when compared to the HF group (*p* = 0.053). Moreover, tadalafil treatment augmented the abundance of the cGMP hydrolyzing phosphodiesterases PDE2 and PDE3 (Figure 2E & F). Thus, the increased abundance of PKG, through inhibition of sGC ^38^ and stimulation of PDE5 ^15^, coupled with elevated levels of the cGMP hydrolyzing PDEs 2 & 3 provide a plausible explanation for the unchanged myocardial cGMP content despite chronic tadalafil treatment.

In the next experiments we sought to elucidate the impact of chronic PDE5 inhibition in HF on key regulators of catecholamine signalling in the heart. We found no difference in β_1_ or β_2_ adrenergic receptor abundance in whole heart homogenates following 4-weeks of tachypacing, in HF or following tadalafil treatment (Figures 3A & B). In agreement with our previous findings ^1^, in HF the protein abundance of G protein coupled receptor kinase GRK2 and protein phosphatases PP1 and PP2A was increased (Figure 3C-E). The increase in GRK2, PP1 and PP2A was established by 4-weeks of tachypacing, reaffirming adrenergic dysfunction is established prior to PDE5 inhibitor treatment in this study. Importantly, chronic tadalafil treatment resulted in a reduction in protein abundance for each of these regulators of β-adrenergic signalling to either the same level (GRK2, PP1) or lower (PP2A) than in control hearts thus mechanistically linking PDE5 inhibition with the enhanced catecholamine responses noted *in vivo*.

**Figure 3.**
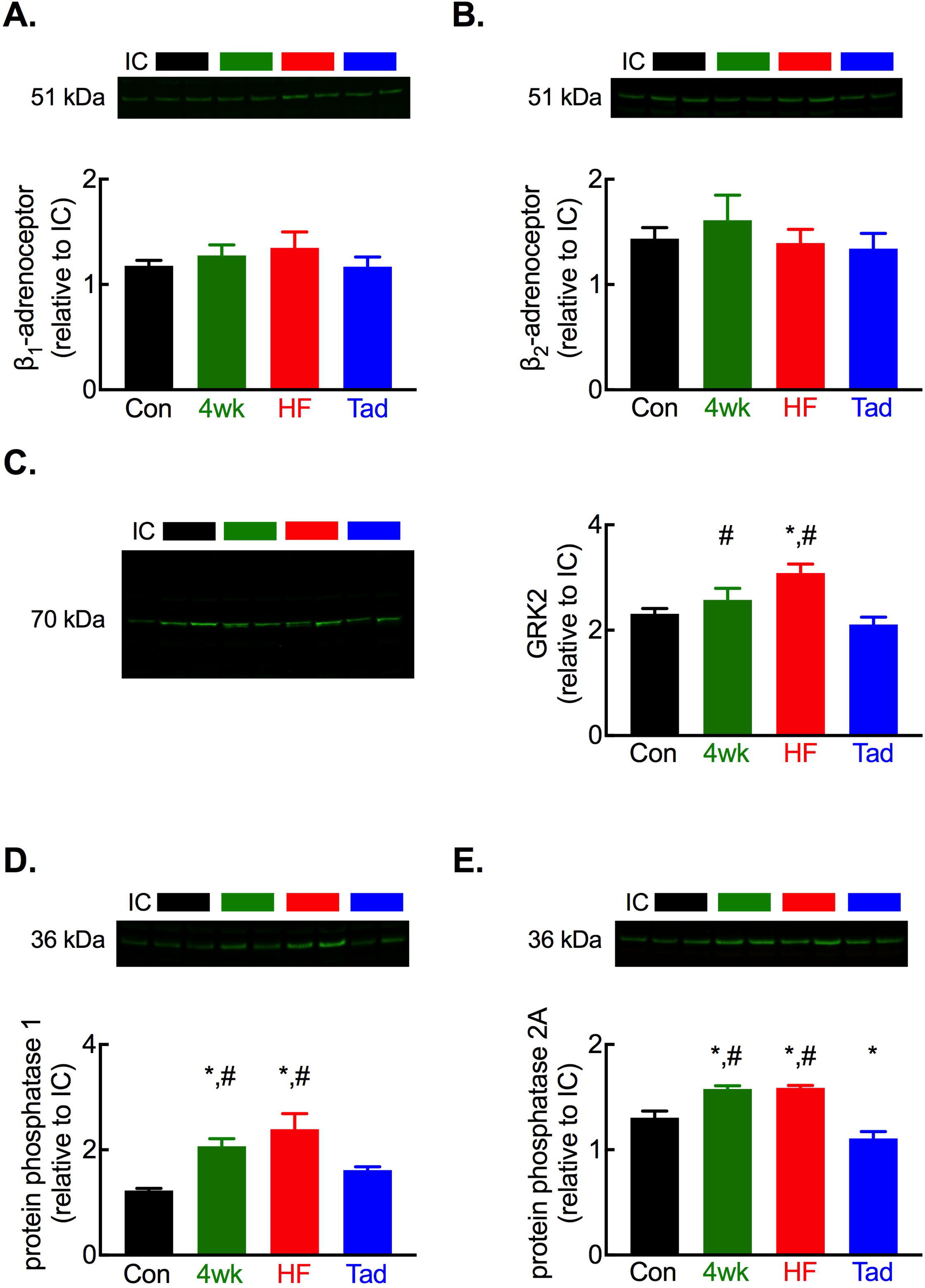
Tadalafil normalizes the abundance of key regulators of β-adrenergic signalling. **A**. Representative blot (upper) and summary data (lower) showing no change in β_1_ adrenergic receptor abundance with tachypacing or tadalafil treatment. **B**. Representative blot (upper) and summary data (lower) showing no change in β_2_ adrenergic receptor abundance with tachypacing or tadalafil treatment. **C**. Representative blot (left) and summary data (right) showing tachypacing induced increased and tadalafil mediated normalisation of GRK2 protein abundance. **D**. Representative blot (upper) and summary data (lower) showing tachypacing induced increased and tadalafil mediated normalisation of PP1 protein abundance. **E** Representative blot (upper) and summary data (lower) showing tachypacing induced increased and tadalafil mediated normalisation of PP2A protein abundance. For all panels *, *p* < 0.05 vs. control; #, *p* < 0.05 vs. tadalafil. *N* = 8 per group.

### PDE5 inhibition restores the transverse tubule network in HF

Since the β_1_-AR and β_2_-AR are functionally distributed on the TT membrane and surface sarcolemma ^5,7^ we sought to determine if the restored response to the β-AR agonist dobutamine following tadalafil treatment was associated with changes in TT density or TT organization in ventricular myocytes (Figure 4A). To quantify changes in TT density we utilized two approaches; i) determination of the distance 50 % of voxels (3 dimensional pixels) are from their nearest membrane (TT or surface membrane) in all three imaging planes, termed ‘half-distance’ ^8,32^ (Figure 4A.b.) and, ii) determination of the fractional area of the cytosol occupied by TT’s in the imaging plane, termed ‘fractional area’ ^8,32^. In agreement with our previous work ^8^ and others e.g. ^9,10,39^ HF resulted in a decrease in ventricular TT density of 19 ± 5 % measured using the half distance (Figure 2B; *p* < 0.001) and 26 ± 5 % using fractional area methodologies (Figure 4C; *p* < 0.001). Importantly, these changes in TT density had occurred by 4-weeks of tachypacing and therefore the TT density reduction observed in HF was established before commencement of tadalafil treatment (Figure 4B-C). Following 3 weeks of tadalafil treatment, with continued tachypacing, there was a complete restoration of TT density back to control levels (half-distance: control, 0.377 ± 0.01 µm; tadalafil, 0.369 ± 0.001 µm; fractional area: control, 0.198 ± 0.01; tadalafil, 0.231 ± 0.01). Despite TT density being fully restored following tadalafil treatment, closer inspection of the skeletonized images (Figure 4A.c.) showed that the TTs are less organized in the tadalafil treated animals than in control myocytes. In control cells TTs are predominantly organized perpendicular to the long axis of the cell (Figure 4D) having a transverse (perpendicularly arranged) to longitudinal (horizontally arranged) ratio of 2.4 ± 0.2 (Figure 4E). After 4-weeks of tachypacing, in end-stage HF and following tadalafil treatment TTs are nearly equally oriented longitudinally and transversely (4-weeks, 1.1 ± 0.1; HF, 1.0 ± 0.1; tadalafil 1.2 ± 0.1; all *p* < 0.001 versus control). Thus, the restoration of TT density following tadalafil treatment is not matched by normalization of their orientation.

**Figure 4.**
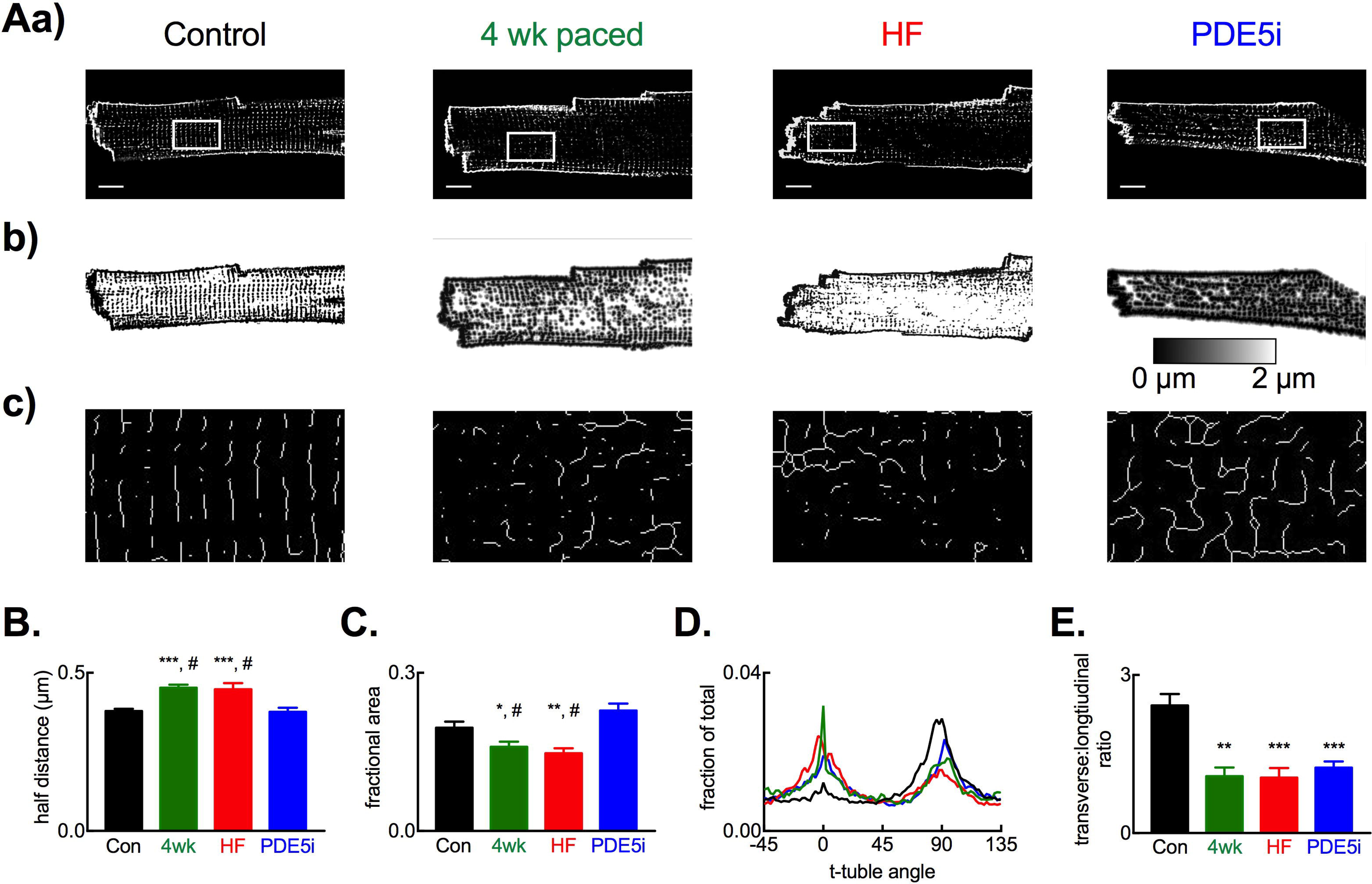
Recovery of transverse tubule density with altered orientation following tadalafil treatment. **A**. Representative images from cell types indicated showing membrane staining (a), distance maps (b) and skeletonized images (c) from regions of interest shown in panel a. Note the reduction in TT density (a, b) and lateralization of TTs in 4-week paced, HF and tadalafil treated groups (c). Scale bars, 10 µm. **B**. Summary data for TT half-distance measurements. **C**. Summary data for TT fractional area measurements. For B and C: n cells / N hearts; control, 30 / 6; 4-weeks, 25 / 4; HF, 29 / 6; tadalafil, 71 / 6. **D**. Representative orientation analysis for TTs in regions highlighted relative to long-axis of cells. 0° represents longitudinally orientated and 90° transversely oriented tubules. **E**. Summary data showing TT transverse to longitudinal orientation ratio. In B, C and E: *, *p* < 0.05 vs control; **, *p* < 0.01 vs. control; ***, *p* < 0.001 vs. control; ^#^, *p* < 0.05 vs tadalafil; by linear mixed models analysis. Panels B, C & E show mean ± SEM.

### Potential molecular mechanisms responsible for changes in TT density in HF and reversal by PDE5i treatment

Although the control of TT turnover remains poorly understood we next sought to identify potential molecular mechanisms responsible for the restoration of TTs following tadalafil treatment. We examined protein abundance of four putative regulators of TT turnover: AmpII (BIN1), junctophilin 2 (JPH2), titin capping protein (Tcap) and myotubularin 1 (MTM1). Of these, JPH2 (Figure 5A) and Tcap (Figure 5B) showed no change in abundance with disease duration or tadalafil treatment whereas MTM1 abundance was increased equally above control in the 4-week tachypaced, HF and tadalafil treated animals (Figure 5C, all *p* < 0.05). Conversely, AmpII protein abundance was reduced from control levels in both the 4-week tachypaced and HF groups where TT density is reduced (Figure 5D.a; *p* < 0.05) and restored to control levels in the tadalafil treated group in line with the restoration of TT density. Moreover, AmpII abundance was also greater in the tadalafil group than both the 4-week tachypaced and HF groups (*p* < 0.05). The relationship between TT density and AmpII protein abundance is examined in Figure 5D.b-c. Here, due to the unpaired nature of the protein samples used for Western blotting and cells used for TT density determination we have used a non-replacement random sampling and fitting simulation approach (1000 iterations) to test the correlation between TT density and AmpII protein abundance. As we have shown previously using gene silencing approaches in isolated myocytes^8^, TT density is correlated with AmpII protein abundance during the development of HF and following treatment with tadalafil (*p* < 0.001).

**Figure 5.**
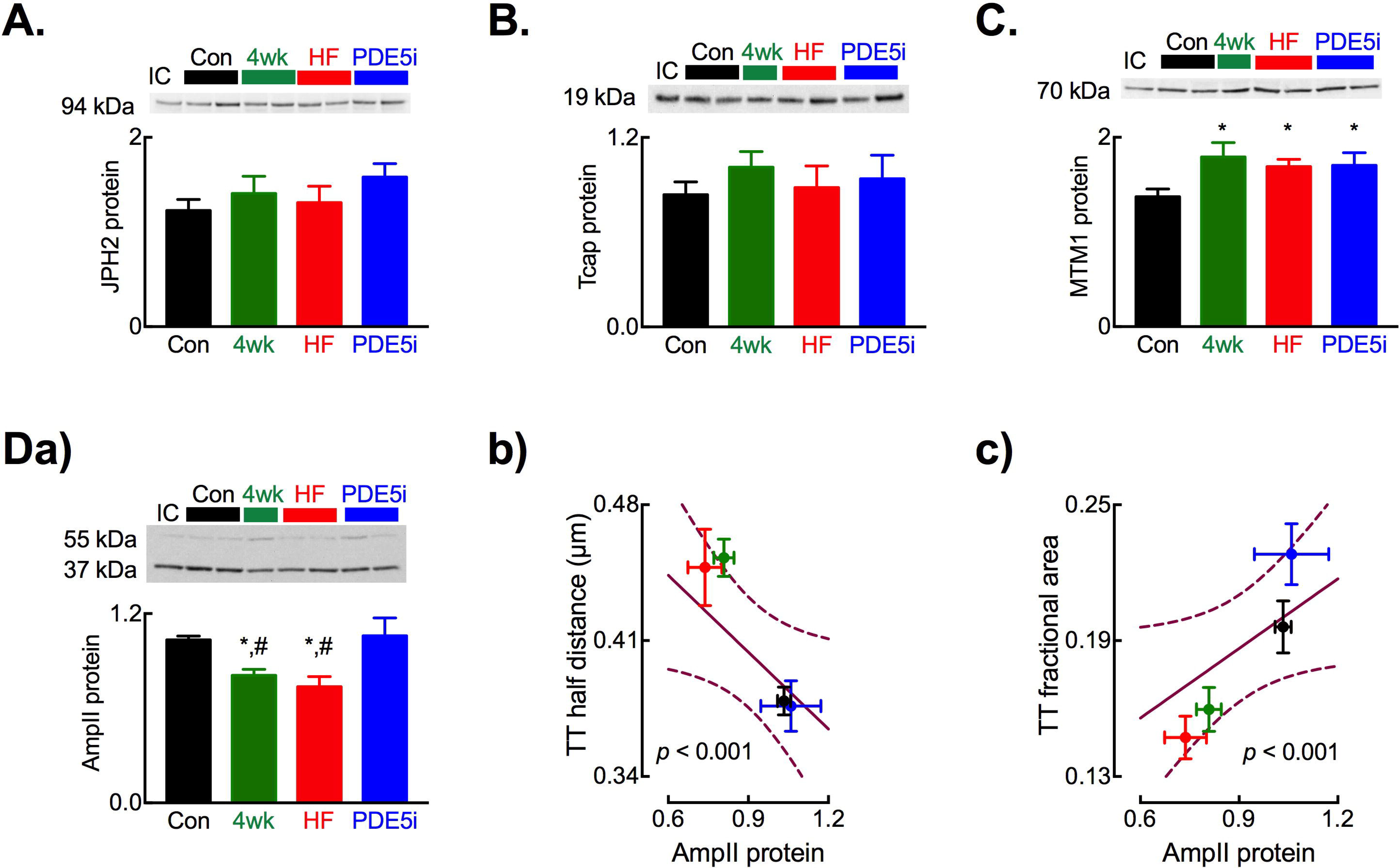
Altered protein abundance of putative regulators of cardiac transverse tubule formation. Representative immunoblots (upper) and summary histograms (lower) for: **A**. JPH2; **B**. TCAP and **C**. MTM1. **D**. Changes in AmpII protein abundance showing (upper) representative AmpII immunoblot and (lower) summary data (a); correlation between AmpII protein abundance and TT half distance (b) and TT fractional area (c). Solid lines through data in b. and c. are linear regressions with slopes significantly different from zero (*, *p* < 0.05). For immunoblots (A – D) *N* = control, 7; 4-weeks, 6; HF, 7; tadalafil, 8 and mean of 3 technical replicates. *, *p* < 0.05 vs. control; ^#^, *p* < 0.05 vs. tadalafil; by linear mixed models analysis. For regression analysis: *n* cells / *N* hearts; control, 30/ 6; 4-weeks, 25 / 4; HF, 29 / 6; tadalafil, 71 / 6. All data presented as mean ± SEM.

Whilst our present data (e.g. Figure 5D) and previous studies ^8,40^ have shown a correlation between AmpII abundance and TT density, there remains some uncertainty over which isoforms of AmpII (BIN1) are expressed in cardiac muscle and drive the formation of TTs in cardiac myocytes. We therefore sought firstly to determine which AmpII isoforms are present within the ovine myocardium. Using RT-PCR approaches and amplifying the variably expressed regions between exons 6 and 18 of AmpII (Supplemental Figure 3) we identified 3 major amplicons in the ovine myocardium (Fig 6A). From amplicon size and subsequent sequence verification we identified the ‘skeletal’, exon 11 containing v8 isoform as the major AmpII isoform expressed in the myocardium with lower levels of expression of v4 and v10 isoforms. By investigating the role of variant 8 AmpII overexpression in NRVMs and iPSC CMs we next sought to establish if the augmented AmpII protein abundance provides a mechanism for the increase in TT density following tadalafil treatment. Importantly, and in contrast to the heterozyous knockout model used by Hong *et al* ^41^, both cell types lack TTs. Both non-transfected and mKate2 fluorescent protein transfected NRVMs have low endogenous AmpII protein abundance whereas AmpII-AC-mKate2 transfection increased AmpII protein abundance 4.6 ± 0.3-fold (Figure 6B, *p* < 0.05). When transfected NRVMs were imaged confocally (Figure 6C), those that were transfected with the mKate2 control vector exhibited diffuse cytosolic and nuclear fluorescence. Conversely, cells transfected with the AmpII-AC-mKate2 vector exhibited widespread intracellular fluorescence in the form of an inter-connected network. To determine if the AmpII driven network formed tubules connected to the surface sarcolemma cells were superfused with the extracellular fluorescent marker Oregon Green 488N. It is clear from Figure 6C that Oregon Green 488N enters the cell and co-localizes with mKate2 fluorescence indicating: i) that AmpII forms tubular structures and, ii) these are patent at and connected to the cell surface. On average 97.9 ± 2.1 % cells transfected with AmpII-AC-mKate2 developed a tubular network compared to no tubules in non-transfected or cells transfected with the mKate2 control vector (Figure 6D, *p* < 0.001). The summary data of Figure 6E demonstrates that AmpII-C-mKate2 expression in NRVMs lead to a robust increase in the fractional area of cells occupied by TTs (59 ± 3 fold, *p* < 0.001). Finally, we also confirmed that AmpII was capable of driving *de novo* tubule formation in human iPSC CMs and that these cells also normally lack a discernible tubular network (Figure 6F-H).

**Figure 6.**
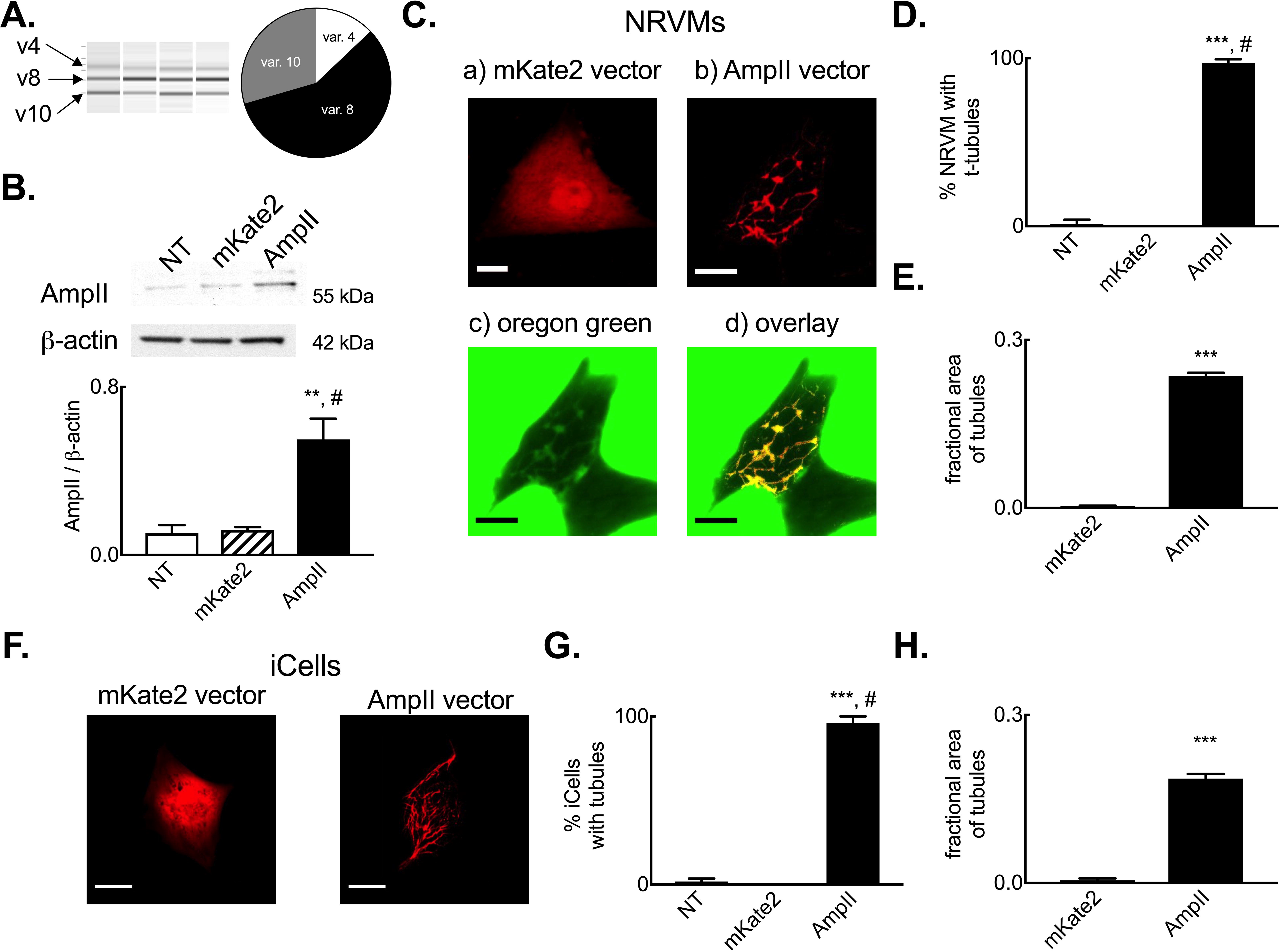
Amphiphysin II drives de novo tubule formation in neonatal rat ventricular myocytes and iPSC derived cardiac myocytes. **A**. Assessment of sheep myocardial AmpII isoform expression showing (left to right) representative Qiaxcel run and summary data on sheep cardiac isoform abundance. *N* = 4 hearts. **B**. Representative immunoblot (upper) and summary data (lower) showing up-regulation of AmpII protein abundance following transient transfection of NRVMs. *N* = 2 isolations. **C**. Representative images of NRVMs transfected with mKate2 control (a) vector or AmpII expression vector (b-d). Extracellular Oregon Green 488 imaging showing a transfected and non-transfected (NT) cell (c) and overlay of the mKate2 channel and OG488N channel (from c) showing co-localisation of markers and patency of AmpII driven tubules only in the transfected cell (d). **D**. Mean data summarizing percentage of non-transfected (NT), mKate2 control vector and AmpII expression vector transfected cells with tubule structures. *N* = 6 isolations (non-transfected, 67 cells; mKate, 23 cells; AmpII, 37 cells) cells in each group. **E**. Mean data summarising fractional area of cells occupied by tubules in mKate2 control vector and AmpII expression vector transfected cells. *N* = 5 isolations (mKate2, 20 cells; AmpII, 29 cells). **F**. Representative images showing AmpII driven tubule formation in iCell iPSC derived cardiac myocytes. **G**. Mean data summarizing percentage of non-transfected (NT), mKate2 control vector and AmpII expression vector transfected iCell iPSC cardiac myocytes with tubule structures (*N* = 2 cell batches; non-transfected, 45 cells; mKate2, 17 cells; AmpII, 40 cells) **H**. Mean data summarizing fractional area occupied by tubules in iCell cardiac myocytes transfected with mKate2 or AmpII vectors. *N* = 2 cell batches (makte2, 17 cells; AmpII, 29 cells). **, *p* < 0.01 vs mKate2; ***, *p* < 0.001 vs mKate2; ^#^, *p* < 0.05 vs NT; by linear mixed models analysis. Scale bars, 10 µm. Panels A, C, D, F & G show mean ± SEM.

## Discussion

The main findings of the present study investigating the impact of chronic PDE5 inhibition on cardiac function and structure in HF are threefold: i) PDE5 inhibition improves indices of cardiac contractility and restores systolic calcium and catecholamine responsiveness; ii) t-tubule density is reduced in HF and fully restored to control levels by PDE5 inhibition and, iii) changes in expression of the putative t-tubule regulator AmpII are correlated with changes in t-tubule density and AmpII drives *de novo* tubule formation in ventricular myocytes. Importantly, in this study we demonstrate that PDE5 inhibition *reverses* rather than *prevents* the HF-dependent changes in cellular structure and catecholamine responsiveness as both the reduction in TT density and loss of catecholamine responsiveness are already present before PDE5 inhibition is commenced.

### PDE5 inhibition mediated reverse remodeling in HF

In the present study tadalafil treatment reversed the tachypacing induced changes in myocardial B-type natriuretic peptide mRNA levels. Consistent with the reversal of B-type natriuretic peptide mRNA we also noted that tadalafil treatment prevented development of a number of subjective signs of HF. The delayed administration of PDE5 inhibition used here distinguishes this study from the majority of previous studies investigating PDE5 inhibition in HF where treatment was initiated simultaneously with, or prior to, disease initiation ^22–27^. Additionally, where earlier studies have implemented delayed PDE5 inhibitor treatment, these appear to have been at a far less advanced stage of HF than in this study based on contractility measurements or the maintained response to catecholamine stimulation ^28–30,37^. Thus, this study adds to a growing weight of evidence on the effectiveness of PDE5 inhibition in treating systolic HF and demonstrates reversal of key elements of the disease process that are evident when cardiac dysfunction is at an advanced stage prior to initiation of PDE5 inhibition.

Whilst changes in B-type natriuretic peptide mRNA are observed, the PDE5 inhibition mediated effects on HF progression occur in the absence of changes in PDE5A mRNA and cGMP levels within the myocardium. Previous studies have also noted no change in PDE5A abundance in preclinical HF models ^29,37^ and Hiemstra *et al*, using a porcine model, reported no change in myocardial cGMP content following tadalafil treatment ^37^. In the present study we investigated potential mechanisms that could explain the lack of change in the global cGMP content of the myocardium and found that PKG (monomer, dimer and total levels) and the cGMP hydrolyzing phosphodiesterases PDE2 and PDE3 were augmented following tadalafil treatment. We suggest therefore that the changes in PKG, PDE2 and PDE3 may underlie the lack of a global change in cGMP, yet, there may still remain sub-cellularly restricted pools of elevated cGMP that may influence distinct signalling pathways and underpin the inotropic, chronotropic and ‘survival’ effects seen with tadalafil treatment.

### PDE5 inhibition in HF: effects on cardiac function and catecholamine responsiveness

The first concern regarding the impact of tadalafil treatment in HF is that of off-target effects. Tadalafil is structurally distinct from other commonly used PDE5 inhibitors (sildenafil, vardenafil) and shows substantially greater (> 1000 fold) specificity for PDE5 over other PDEs than both sildenafil and vardenafil with the exception of PDE11 where the selectivity ratio (PDE5 IC_50_: PDE11 IC_50_) is between 7.1 and 40 42-45. However, the absence of PDE11 in cardiac myocytes ^46^ together with the lack of any obvious neurological disturbance or myalgia (PDE11 dependent side-effects) would suggest that despite the plasma concentration of tadalafil used (~ 50 nmol/l) causing complete inhibition of PDE5 (IC_50_ range 0.94 – 9.4 nmol/l) ^42,47^, the approximate 40 %, inhibition of PDE11 is not underpinning the cardioprotective effects observed in the present study. The development of PDE11 specific antagonists or use of lower doses of tadalafil in future studies would enable this to be conclusively determined.

In those studies where PDE5 inhibition was started at varying time points following disease initiation there is indeed evidence of attenuated deterioration in function or some reversal of extant dysfunction ^28,30,48,49^. However, in each of these cases catecholamine response data and commentary on the development of signs of HF are absent. Conversely, in those studies where the systolic calcium transient or adrenergic responsiveness have been measured they are either maintained ^29^ or augmented ^37^ in the ‘HF’ groups suggesting that the disease stage in these particular studies is relatively early. In comparison, the present study, using a large animal model of tachycardic end-stage HF, demonstrates that contractile and adrenergic dysfunction are established before the onset of PDE5 inhibitor treatment and these effects are at least partially restored by tadalafil treatment.

In the present study we investigated the potential mechanisms underpinning the restoration of catecholamine responsiveness following PDE5 inhibition. Whilst we found no change in β_1_ or β_2_ adrenergic receptor density in HF or following tadalafil treatment it remains possible that the restoration of TT density with tadalafil treatment is important in directing the sub-cellular aspects of β-receptor signaling and thence augmenting cardiac contractility ^5,50^. However, in line with the restoration of catecholamine sensitivity we observed tadalafil dependent decreases in GRK2, PP1 and PP2A protein abundance which would increase cAMP levels ^51^ and enhance target phosphorylation including sites on myofilament proteins, phospholamban and the L-type Ca^2+^ channel ^52–54^.

### PDE5 inhibition in HF: effects on cardiac cellular structure

Increasing evidence implicates a reduction in TT density as a major factor contributing to cardiac dysfunction in diverse cardiac diseases ^8-10,22,32,39,55-57^. In the present study, we also note a decrease in TT density in HF and importantly that this is fully reversed by PDE5 inhibition. Our findings extend the earlier studies of Huang *et al* ^27^ where sildenafil treatment commenced concurrently with aortic-banding prevented TT loss and Xie *et al* ^22^ where delayed sildenafil treatment partially restored right ventricular TT density in a pulmonary artery hypertension model. The incomplete recovery of TTs noted by Xie *et al* may reflect the shorter treatment duration (2 weeks), drug choice (sildenafil) or chamber-wall strain differences (right versus left) ^58–60^.

Whilst tadalafil treatment restores TT density, the TTs in this group are equally distributed in the transverse and longitudinal orientations, rather than predominantly transverse orientation seen in control hearts. Previous studies have noted plasticity in TTs and their ability to be restored to varying extents following interventions to reverse HF ^27,60–62^ although these have not investigated TT orientation in the various intervention arms. Here, we sought to determine if changes in TT density in HF and following tadalafil treatment are correlated with changes in the abundance of proteins implicated in TT biogenesis in striated muscle ^63–65^. Of the putative candidate proteins only one, AmpII, correlates with TT density changes in HF and following tadalafil treatment. AmpII (BIN1) is known to be involved in TT biogenesis in skeletal muscle ^66^ and in cardiac muscle protein abundance is reduced in HF ^8,40,61^. Using siRNA gene silencing in cultured adult ventricular myocytes we have shown that AmpII is required for cardiac TT maintenance ^8^ and Hong *et al* ^41,67^ and more recently, De La Mata *et al* ^68^ demonstrated the importance of BIN1 (AmpII) in delivery of L-type Ca channels to TT membranes and TT membrane folding. In the present study, we extend this earlier work and show that AmpII is capable of driving the *de novo* formation of tubule like structures in both NRVMs and iPSC CMs; model cell types that lack TTs. Importantly, co-labelling with extracellular dye shows that the AmpII driven tubules are patent at the cell surface. Thus, an emerging body of work highlights the importance of AmpII in the TT biology in the heart and future studies will be required to define the functional significance of AmpII driven TT formation.

Moreover, and in stark contrast to the work of Hong *et al* using the mouse ^41^, we also demonstrate that exon 11 (previously known as exon 10 ^69^) containing variants 8 and 4 of AmpII (BIN1) are expressed within the sheep myocardium with variant 8 mRNA being the dominantly expressed isoform. We also show that variant 8 is capable of inducing *de novo* tubulation in cardiac cells. Further work will be required to elucidate the significance of species differences in AmpII isoform usage and the critical components of AmpII for cardiac muscle TT formation; it may be that, as with chamber differences in TT density, important distinctions exist between laboratory rodents and large mammals including human ^8,32,70^.

Beyond the correlation between AmpII protein abundance and TT density, we also noted a generalized increase in lipid phosphatase MTM1 protein abundance in the 4-week tachypaced, HF and tadalafil treated animals. In skeletal muscle mutations in MTM1 or altered interaction between MTM1 and a number of binding partners, including AmpII lead to inherited centronuclear myopathies, some of which are associated with cardiomyopathies ^71,72^. Furthermore, MTM1 augments AmpII driven membrane tabulation ^72^ and MTM1 deficiency leads to reductions in skeletal muscle TT density ^73,74^. Thus, it is possible the increases in MTM1 protein abundance noted in the present study is a compensatory mechanism to maintain TTs in the face of cardiac pathology. However, whether the increase in MTM1 abundance is responsible for the lateralization of TTs in HF and tadalafil treated animals remains to be determined although addressing this in the iPSC and NRVM model systems would be problematic given the inherent disorder of the AmpII driven tubules in these cell types. Finally, there remain the unanswered questions as to whether: i) TT loss is a cause or consequence of HF and, ii) if tadalafil improves cardiac function and this allows TT density to recover or, tadalafil drives TT recovery and thence improves cardiac function. Irrespective of these potential caveats, the dependence of TT maintenance on AmpII ^8^ and the demonstration in the present study that AmpII drives *de novo* tubule formation in cardiac myocytes highlights AmpII as a target for future studies. Selectively manipulating AmpII protein abundance in established and evolving cardiac disease states in future studies could resolve this question.

### Limitations

There are some limitations to the present study that arise primarily from the lack of suitable reagents exhibiting satisfactory cross-reactivity with sheep. As such, we have been unable to serially assess natriurtetic peptide (BNP or NT-proBNP) levels in plasma to monitor the progression and resolution of HF; however, we do observe a strong up-regulation of myocardial NPPB (BNP) mRNA in HF and normalization in the tadalafil treated animals using a single time point assay in each group of experimental animals. The main operator was not blinded to treatment and the presence of signs of HF (lethargy, dyspnea etc) are subjectively assessed; critically however, the changes in myocardial NPPB (BNP) mRNA validate the qualitative assessments regarding symptom free survival. Additionally, we have been unable to determine PDE5A protein abundance and have relied on quantitative PCR based assessments. Whilst our work shows no change in the myocardial cGMP content we cannot rule out the possibility that cGMP is elevated in certain sub-cellular pools and thus mediates the effects of tadalafil treatment that we observe. FRET based sensors may offer some potential to assess sub-cellular cyclic nucleotide pools, their use in sheep would require maintenance of primary cardiac myocytes in extended culture to allow transgene expression and thus significant concerns would arise from cellular de-differentiation and ultrastructural changes that inevitably occur when adult cardiac cells are maintained in culture conditions. A further limitation in the present study arises from the anatomical alignment over the cardiac apex over the sternum in adult sheep and inaccessibility to a suitable imaging window to allow four-chamber apical views for a thorough echocardiographic assessment of ejection fraction and E/A ratios. Furthermore, we have found cardiac conductance catheterization to be of limited use given interference from the pacing lead in the right ventricle. Nevertheless, short axis trans-thoracic imaging is possible and clearly shows marked cardiac dilatation and reduced contractility in HF that is at least partially normalized by tadalafil treatment.

In summary, using a large animal model of end-stage HF we have shown that chronic treatment with the PDE5 inhibitor tadalafil reverses already established cellular ultra-structural remodeling and impaired catecholamine responses and also restores systolic calcium in HF. Given the localization of β-ARs and G-protein coupled regulators of cardiac catecholamine responses across the TT and surface sarcolemma, the recovery of TTs following PDE5 inhibition provides a cellular mechanism for the restoration of catecholamine reserve in tadalafil treated animals.

## Methods

### Ethical Approval

All experiments were conducted in accordance with The United Kingdom Animals (Scientific Procedures) Act, 1986 and European Union Directive EU/2010/63. Local ethical approval was obtained from The University of Manchester Animal Welfare and Ethical Review Board. Reporting of animal experiments is in accordance with The ARRIVE guidelines ^75^.

### Ovine HF model and drug treatment

HF was induced in 69 female Welsh Mountain sheep (~ 18 months) by right ventricular tachypacing as described previously ^1,8,32-34,76,77^. In brief, anesthesia was induced and maintained by isoflurane inhalation (1 – 5 % in oxygen). Under aseptic conditions using transvenous approaches a single bipolar endocardial pacing lead was actively fixed to the right ventricular apical endocardium and connected to a cardiac pacemaker (e.g. Medtronic Sensia, Medtronic Inc. USA) and buried subcutaneously in the right pre-scapular region. Peri-operative analgesia (meloxicam, 0.5 mg/kg) and antibiosis (enrofloxacin 5mg/kg or oxytetracycline 20mg/kg) were administered subcutaneously and animals allowed to recover post-operatively for at least 1 week prior to commencement of tachypacing (210 beats per minute; bpm). Animals were monitored at least once daily for onset of clinical signs of HF including lethargy, dyspnea and weight loss. Following 28 days of tachypacing animals (*N* = 27) were randomly assigned to receive the PDE5i tadalafil. Treated animals were administered 20 mg tadalafil (Cialis; Lilly, Netherlands) orally once daily for a further 3 weeks and tachypacing (210 bpm) was continued throughout the treatment period. The designated endpoint for the study was either the onset of signs of HF (dyspnea, lethargy, weight loss) or 49 days of tachypacing (21 days tadalafil treatment).

### *In vivo* assessments of cardiac contractility and catecholamine responsiveness

During echocardiographic and electrocardiographic assessments tachypacing was stopped for 15 minutes prior to data acquisition. Following *in vivo* functional assessments tachypacing was continued at 210 bpm.

Cardiac dimensions and contractility were assessed by right para-sternal transthoracic echocardiography (Vivid 7; GE Healthcare, UK) in non-sedated conscious sheep gently restrained in a supine position. Due to the anatomical alignment of the ventricular apex over the sternum in sheep we are unable to obtain satisfactory 4-chamber long axis views and therefore calculate ejection fraction. However, fractional shortening was measured using m-mode imaging at a level proximal to the papillary muscles and fractional area change calculated from short-axis 2d views at the mid-papillary level ^31^.

Heart rate was determined from a five-lead ECG (Iox, EMKA Technologies, France). The chronotropic response to β-adrenergic stimulation was assessed as described previously ^31^ by intravenous infusion of dobutamine hydrochloride (0.5 – 20 µg/kg/min) prepared in 0.9 % saline (Baxter, UK) and delivered using an infusion pump (Harvard Apparatus, UK). Heart rate was determined after 5 mins of dobutamine infusion at each dose and then the infusion rate was doubled to a maximum dose of 20 µg/kg/min or until the heart rate reached 240 bpm.

### Transverse tubule staining and quantification

Single left ventricular mid-myocardial myocytes were isolated using a collagenase and protease digestion technique as described in detail previously ^1,8,32,77^. Briefly, animals were killed by an overdose of pentobarbitone (200 mg/kg intravenously). Heparin (10,000 units) was also used to prevent coagulation. The heart was rapidly excised and left anterior descending coronary artery cannulated and perfused with collagenase (Worthington Type II, 0.1mg/ml) and protease (from *Streptomyces griseus*, 0.02 mg/ml) containing solutions and single myocytes dispersed by gentle trituration in a taurine solution containing (in mmol/l): NaCl, 113; taurine, 50; glucose, 11; HEPES, 10; BDM, 10; KCl, 4; MgSO_4_, 1.2; Na_2_HPO_4_, 1.2; CaCl_2_, 0.1 and BSA, 0.5mg.ml^-1^ (pH 7.34 with NaOH). Changes in intracellular Ca^2+^ concentration were measured using Fura-2 at 37 °C in cells electrically stimulated at 0.5 Hz as we have described previously ^1,33,78^.

TT’s were visualized by staining with di-4-ANEPPS (4 µmol/l. Invitrogen, UK) as described in detail previously ^8,32,70^. Stained myocytes were imaged by confocal microscopy (Leica SP2 or Zeiss 7Live; 488 nm excitation and 505 nm long pass emission settings) at 100 nm XY pixel dimensions and 162 nm Z-step size. Images were deconvolved using the microscope specific derived point spread function (PSF) using Huygens Software (Scientific Volume Imaging, Netherlands)^8,32,70,79^. Following background correction and image thresholding the following quantitative assessments of TT’s were calculated: i) TT fractional area derived from binarized images as the pixels occupied by TT’s as a fraction of the planar cell area (ImageJ. NIH, USA), ii) the distance 50 % of voxels in the cell are from the nearest membrane (TT or surface sarcolemma), hereafter referred to as the half-distance, was determined using custom algorithms written in IDL (Exelisvis, UK) ^8,32,70^. Images were first interpolated to give the same X, Y, Z spacing (‘congrid’) and then Euclidean distance maps determined (‘morph_distance’) and, iii) TT orientation was determined from skeletonized binary images as described previously ^8,57^ using the ImageJ plugin ‘Directionality’.

### mRNA, protein expression and cGMP content analysis

Changes in protein expression were assessed in samples using approaches described in detail previously ^1,8,33^. Following cardiac excision, a ~ 1 cm^3^ region of the posterior left ventricular free wall was removed and snap frozen and stored in liquid nitrogen until analysis. Whole myocardium homogenates (~ 100 mg starting material) were prepared in RIPA buffer with protease and phosphatase inhibitors (0.1 mg/ml phenylmethanesulphonylfluroide, 100 mmol/l sodium orhtovanadate, 1mg/ml aprotonin, 1mg/ml leupeptin) and protein content determined (DC Protein Assay, BioRad, UK). Proteins were separated by PAGE, transferred to nitrocellulose membranes and protein expression determined using the antibodies and blotting conditions outlined in Table 2. For PKG protein measurements, sample preparation and blots were conducted under non-reducing conditions. Membranes were visualized by chemiluminescence (Syngene, UK) or IR-Dye labeled secondary antibodies (Licor, UK). Previously we determined that the ‘classical’ housekeeping proteins GAPDH and β-actin have altered expression in various disease states and therefore we normalized expression of our protein of interest to an internal control (IC) common to each blot *and* total protein transferred to the membrane determined *either* by Ponceau S staining or REVERT total protein stain (Licor, UK). The IC was derived from a single control sheep (which itself was not included in the present study). Blots were repeated in triplicate on separate occasions and data averaged to minimize any effects due to pipetting or protein transfer errors. Full length blots and validation controls are shown in Supplemental Information (Figure S1).

**Table 2.**
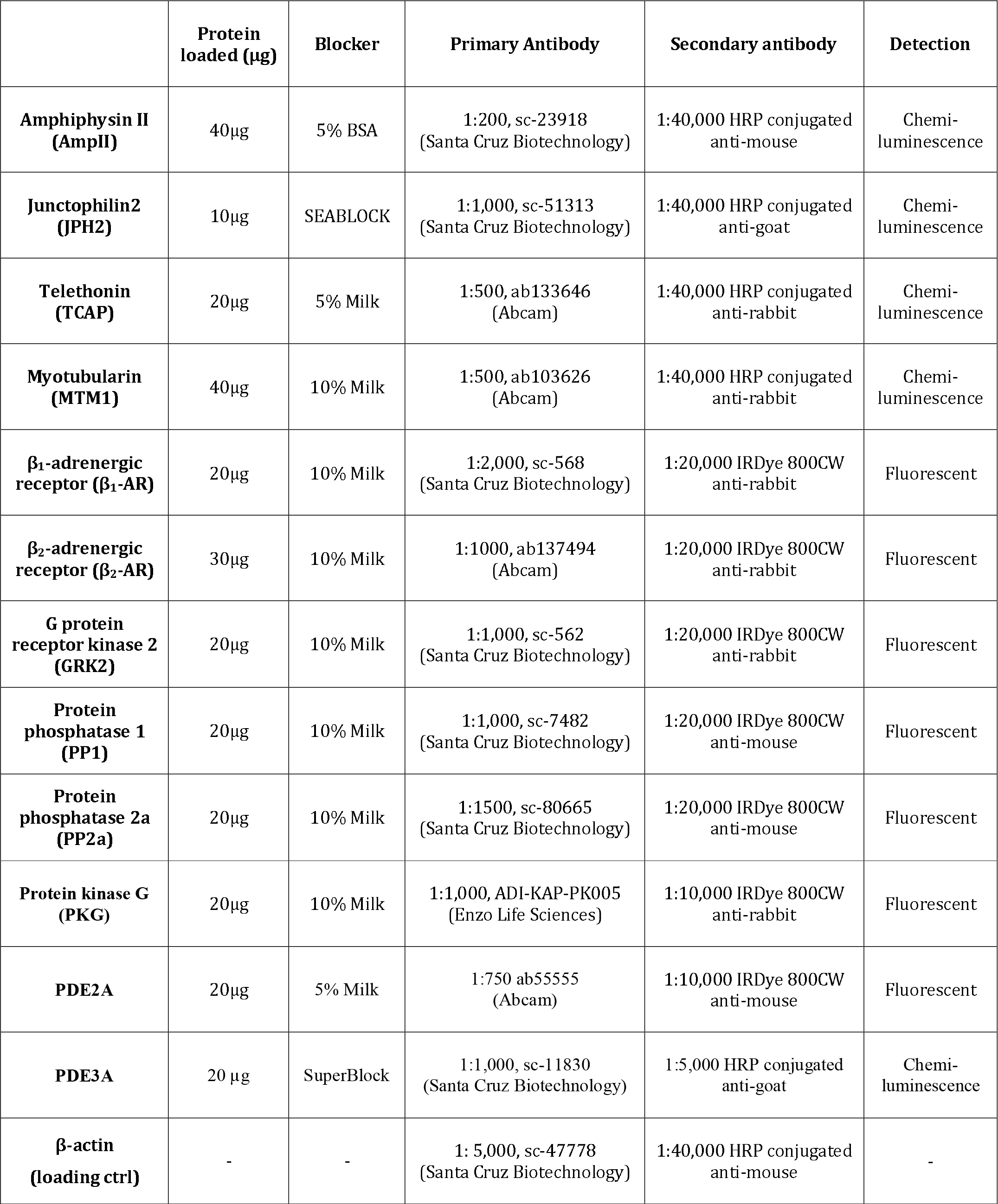
Blotting conditions for selected proteins

Total RNA was isolated using Trizol reagent, purified using RNAeasy columns (Qiagen, UK) following manufacturers recommendations. RNA yield and integrity were determined using a Nanodrop (ThermoFisher, UK) and TapeStation (Agilent, UK) instruments respectively. 1 µg total RNA (RIN scores > 7.0) was then reverse transcribed using a high capacity RNA to cDNA kit (Applied Biosystems, UK) following manufacturers steps. PDE5A (Oa04859755_m1) and NPPB (Oa04931155_g1) mRNA abundance was determined using Taqman single tube FAM-MGB assays (ThermoFisher, UK) following the manufacturers protocols and normalized to either RPLPO alone (Oa04824512_g1) or TBP (Oa0481075_m1) and YWHAZ (Oa03216375_gH) combined using the 2^−ΔΔC^ method (Supplemental Information, Figure 2). All qPCR experiments were repeated 3 times and data averaged. For AmpII (BIN1) isoform determination the variably expressed region between exon 6 and exon 18 (Supplemental Information, Figure 3) was assessed using standard PCR approaches using primers (Eurofins, Germany; forward) 5’-TACGAGTCCCTTCAAACCGC, (reverse) 5’-AGTGTCAACGGCTCTTCCAG. Phusion high fidelity DNA polymerase with GC rich buffer was used for hot start PCR amplification (98 °C, 30 sec; 40 cycles; 98°C - 10sec, 62.1 °C - 30 sec, 72 °C - 30 sec; 72 °C - 10 min) and amplicons separated using a Qiaxcel Advanced Separation System using 15 - 3000 bp markers (Qiagen, UK).

Myocardial cGMP content was determined by competitive ELISA (Amersham, UK) following the manufacturers non-acetylation procedure. Tissue was homogenized in 6 % (w/v) trichloracetic acid, centrifuged and the supernatant washed 4 times before lyophilisation under nitrogen at 60 °C and proceeding to the immunoassay.

### Isolation of Neonatal Rat Ventricular Myocyte, induced pluripotent stem cell cardiac myocyte culture and AmpII overexpression

Neonatal rat ventricular myocytes (NRVMs) were isolated from 2-day old Wistar rats and plated onto 8 well plates (Ibidi µ-plates; Thistle Scientific UK) at a density of ~ 3 × 10^5^ cells/ml in media (containing: DMEM, 68 %, M199, 17 %; normal horse serum, 10 %; fetal bovine serum, 5 %; all Gibco Life Technologies, UK). Fungizone, Penicillin (10,000 units/ml) and streptomycin (10 mg/ml) were added to a final concentration of 1 % and bromodeoxyuridone (100 µmol/l) added to retard fibroblast proliferation. Cells maintained at 37 °C in a 5 % CO_2_ incubator and media changed every 2 – 3 days. In brief, rat pups were killed by cervical dislocation, bodies rinsed in 70 % ethanol and the ventricles rapidly excised and placed in ice cold dissociation buffer (mmol/l: NaCl, 116; HEPES, 20; glucose, 5.6; KCl, 5.4; NaH_2_PO_4_, 1; MgSO_4_, 0.83; pH 7.35). Single NRVMs were digested using 0.75 mg/mL collagenase A (Roche, UK) and 1.3 mg/ml pancreatin (Sigma, UK) by gentle stirring (120 rpm, 7 minutes, 37 °C) and trituration with a 25 ml Stripette. NRVMs (1 – 3 days post isolation) were transiently transfected with either variant 8 of human BIN1 (AmpII) (Origene Inc, USA) cloned into pCMV6-AC-mKate2 entry vector (Origene Inc, USA) or pCMV6-mKate2 as a negative control. Plasmid DNA (6 µg) was mixed (1 : 3 ratio) with transfection reagent (Fugene 6; Promega, UK) in reduced serum media (OptiMEM; Gibco Life Technologies, UK) and cells transfected and maintained in a 5 % CO_2_ incubator at 37 °C for 48 hours prior to visualization using a Nikon A1R^+^ confocal microscope (excitation, 561 nm; emission 595 ± 50 nm). The density (fractional area) of tubular structures was determined as described above for adult ventricular myocytes.

Human induced pluripotent stem cell derived cardiac myocytes (iPSC CMs) were purchased (Cellular Dynamics International, USA) and stored in liquid nitrogen until use. Cells were thawed, plated (~6 × 10^4^ cells/cm^2^) and maintained as per the manufacturer’s instructions and using the manufacturers supplied media (iCell cardiomyocyte plating and culture media). Media was changed every 2 days. Five days after plating cells were transfected as described for NRVMs. Tubular structures were quantified by determining the fractional area as described above.

### Determination of plasma tadalafil concentration and plasma protein binding

Plasma tadalafil concentration (*N* = 13) was determined by mass spectrometry. Briefly whole blood from chronically treated and tachypaced animals was collected from the jugular vein in to K_2_ EDTA blood collection tubes (Vacutainer; Becton Dickinson & Co, UK) on the day of euthanasia and stored on ice. Following centrifugation (4000 *g* for 15 minutes, 4 °C; within 30 minutes of sample collection) plasma was stored in liquid nitrogen until analysis. Tadalafil concentrations were subsequently determined using a liquid chromatography-mass spectrometry method modified from Kim *et al* ^80^. Briefly, following metabolite extraction samples were spiked with heavy stable-isotope containing Tadalafil and Sildenafil (Toronto Research Chemicals, Canada). Each sample was analysed using a Selected Reaction Monitoring method on a ThermoFisher Accela UHPLC coupled to a ThermoFisher TSQ Vantage, using transitions described by Kim *et al* ^81^. with sample blanks and tadalafil spiked samples analysed periodically to ensure quantitative accuracy. Tadalafil concentration was calculated by comparing the peak area of the endogenous compound to that of its heavy labelled counterpart. Tadalafil plasma protein binding was also determined by mass spectrometry following rapid equilibrium dialysis (ThermoFisher, UK). K_2_EDTA plasma samples (100 µl) from tadalafil treated animals were dialysed in to phosphate buffered saline following manufacturers instructions. Following dialysis equal volumes of control plasma or phosphate buffered saline were added respectively to the dialysate and sample plasma prior to mass spectrometry. The percentage plasma protein bound tadalafil was calculated as (dialysate concentration / sample concentration) × 100.

### Statistics

Data are presented as mean ± standard error of the mean (SEM) for *n* observations / *N* experiments (animals). Where multiple observations (*n*) have been obtained from the same animal (*N*) linear mixed modeling (SPSS Statistics. IBM, USA) was performed thus accounting for the nested (clustered) design of the experiment e.g. multiple observations from the same heart and replicate Western blots. Data were log_10_ transformed prior to linear mixed modeling ^82^. Dose response and echocardiographic experiments were assessed using a linear mixed modelling (‘xtmixed’; Stata, StataCorp USA), one-way analysis of variance with Bonferroni correction for multiple comparisons (‘oneway’; Stata, Stacorp USA) or repeated measures ANOVA with Holm-Sidak multiple comparison correction as indicated in figure legends. Survival data and Cox’s proportional hazard ratios were also assessed using Stata (‘stcox’). Linear regressions were fitted to non-paired data (Figure 5) by simulating 1000 randomly sampled (7 points in each condition without replacement) data subsets and fitting each of these with a linear regression. The mean slope and 95 % confidence limits were then obtained and superimposed on the plotted mean ± SEM data points. All values were considered significant when *p* < 0.05.

### Data & Code Availability

The data that support the findings of this study are available from the corresponding author on reasonable request. Similarly, computer code (IDL) used for assessment of TT density are available from the corresponding author on reasonable request.

## Acknowledgements

This work was supported by grants from The British Heart Foundation: FS/12/57/29717, CH200004/12801, FS/15/28/31476, FS/10/71/28563, PG/15/70/31724, IG/15/2/31514, FS/09/002/26487, FS/12/34/29565, FS/15/67/32038, FS/10/52/28678 and Medical Research Council: MR/K500823/1, MR/K501211/1. The authors also acknowledge technical support from Medtronic UK. and Oscor Inc. (USA) in the form of high rate pacing patches for pacing devices and adaptors for pacing leads respectively.

## Author Contributions

Study Conception & Design: AWT; Supervision: AWT, DAE, KMD; Animal models, *in vivo* data generation and analysis: ML, AWT, EJR, CMP, GJK, GWPM, LSW, DCH & LKB; mRNA, protein and cGMP studies CERS, EJR, JLC & RFT; transverse tubule analysis and protein regulator studies, JLC; Mass spectrometry: SJC & RDU. All authors contributed to manuscript drafting / editing.

## Competing Interests

Financial, none; Funding, none; Non-Financial, none.

