## Supplemental Information for "Phosphodiesterase 5 inhibition improves contractile function and restores transverse tubule loss and catecholamine responsiveness in heart failure"

+44 161 275 7969

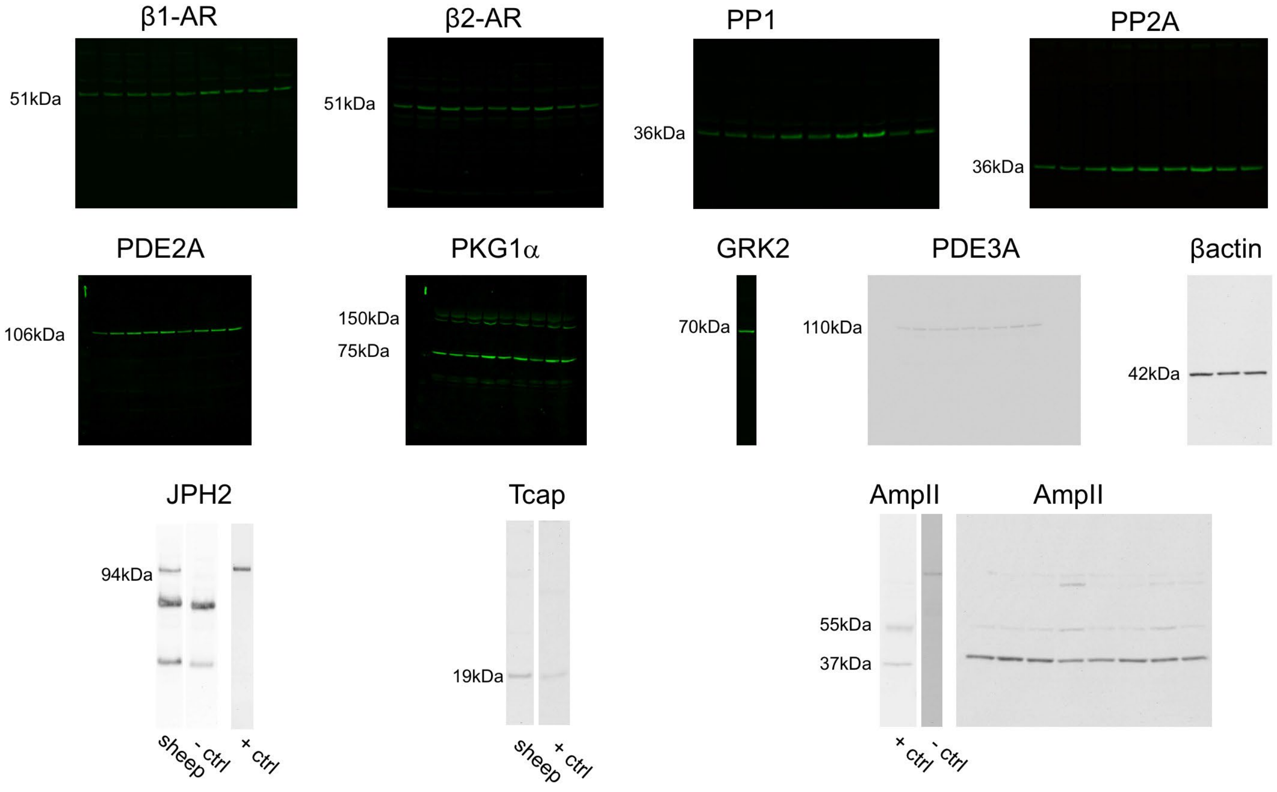


**Supplemental Figure 1. Full length blots.**

Full length blots of proteins of interest obtained using either a fluorescence or chemiluminescence based image capture system. Proteins and molecular weights as indicated. +ctrl denotes positive control (rat foetal heart for Tcap; mouse liver for MTM1 and rat skeletal muscle for AmpII); - ctrl denotes negative control (secondary antibody only).

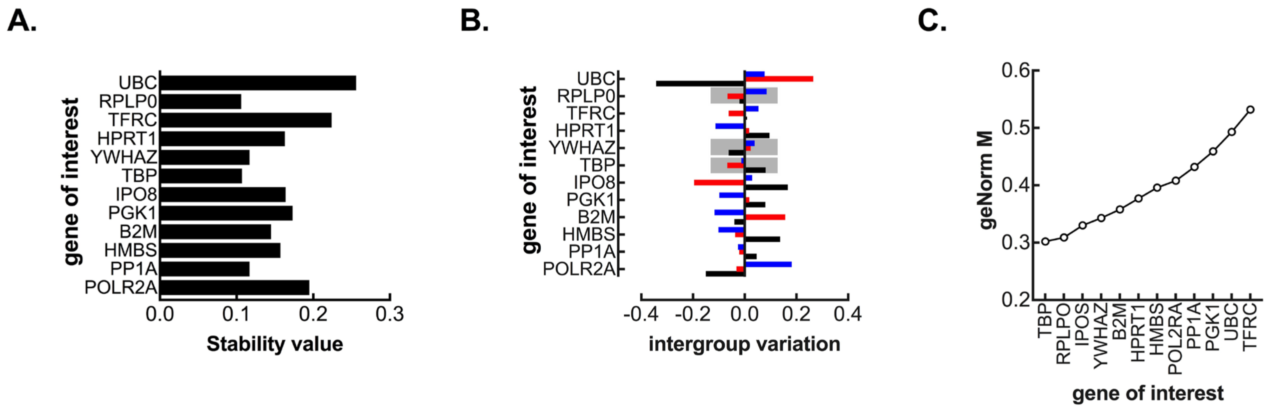


**Supplemental Figure 2: Housekeeping genes used for quantitative PCR normalisation in sheep ventricular samples.**

**A.** NormFinder stability values for genes evaluated as potential normalisation controls. **B.** NormFinder intergroup variation values for genes evaluated as potential normalisation controls (black, control; red, heart failure; blue, tadalafil). The highlighted genes were selected as normalisation controls. **C.** geNorm stability values for genes indicated.

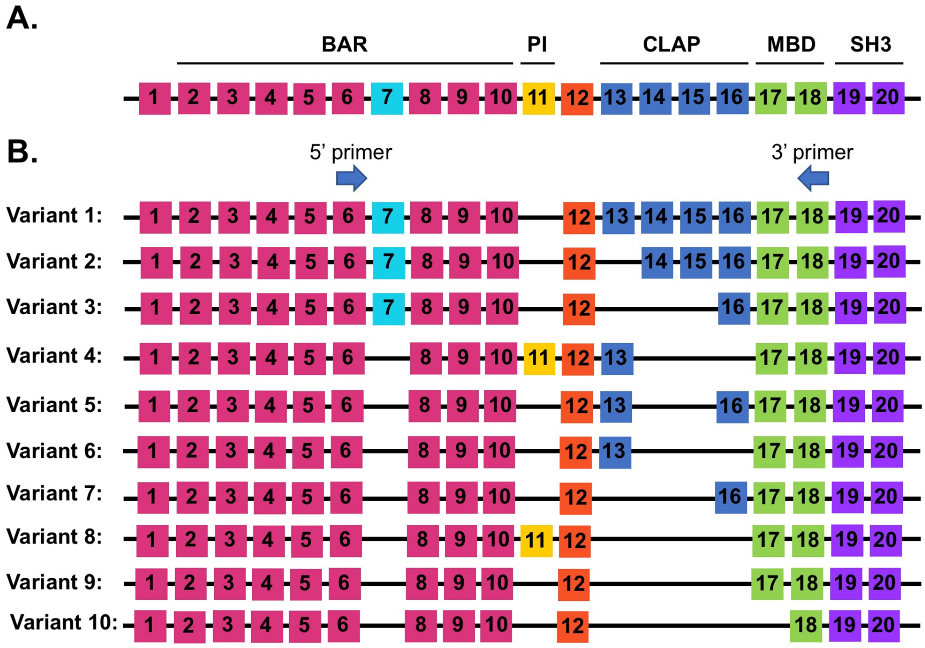


C.

| variant | amplicon (bp) |
|---|---|
| 1 | 1020 |
| 2 | 892 |
| 3 | 762 |
| 4 | 736 |
| 5 | 795 |
| 6 | 691 |
| 7 | 670 |
| 8 | 608 |
| 9 | 563 |
| 10 | 495 |

**Supplemental Figure 3. Primer design strategy for AmpII isoform detection in sheep ventricular myocardium.**

**A.** Exon structure for AmpII gene based on sheep (accession XM_012147686.2) and human (accession NM_139343.2) sequences. BAR, Bin-Amphiphysin-Rvsp domain; PI, phosphoinositide binding domain; CLAP, clathrin-AP2 domain; MBD, myc binding domain; SH3, src homology domain). **B.** PCR primers were designed to span the variable region between exons 6 and 18. **C.** Predicted amplicon sizes for variants 1 – 10 of AmpII.
